# Suppression of corticospinal excitability by sleep spindles without increase in GABAergic inhibition

**DOI:** 10.64898/2026.09.03.748984

**Authors:** Friederike Breuer, Sianna Groesser, Katrin Himmerich, Katrin Knietsch, Umair Hassan, Saman Seifpour, Anastasiia Grigoreva, Ulf Ziemann, Til Ole Bergmann

## Abstract

Thalamocortical sleep spindles are hypothesised to support memory consolidation during sleep by creating transient windows of enhanced hippocampal-neocortical communication and synaptic plasticity. A recent real-time electroencephalography (EEG)-triggered transcranial magnetic stimulation (TMS) study found a pulsed suppression of corticospinal excitability during spindles relative to spindle-free non-rapid eye movement (NREM) sleep, driven by the spindle falling phase. We hypothesised that this phasic suppression may reflect local inhibitory network dynamics, measurable as GABA-A receptor-mediated short-interval intracortical inhibition (SICI) using paired-pulse TMS. We applied real-time EEG-triggered single- and paired-pulse TMS over the primary motor cortex during pre-sleep wakefulness, spindle-free N2/N3 sleep, and at four sleep spindle phases (peak, falling, trough, and rising). Corticospinal excitability was strongly reduced from wakefulness to spindle-free N2/N3 sleep, and further suppressed during sleep spindles. Numerically, excitability was lowest during the falling phase and trough, although we found no significant modulation across spindle phases. Contrary to our hypothesis, neither spindle presence nor phase significantly modulated SICI. Secondary analyses provided preliminary evidence that slow oscillations present at stimulation increased excitability and reduced SICI, irrespective of spindle presence. Together, these findings indicate distinct contributions of sleep/wake vigilance states, sleep spindles, and slow oscillations to cortical network dynamics, and provide new insight into the transient modulation of corticospinal excitability and GABA-A-receptor mediated SICI during human NREM sleep.

## Introduction

Thalamocortical sleep spindles, oscillatory bursts of 10-15 Hz, are one of three cardinal oscillations in non-rapid eye movement (NREM) sleep considered to orchestrate the reactivation of memories during sleep (Fernandez & Lüthi, 2020). According to the active systems consolidation hypothesis, repeated reactivation of initially labile hippocampus-dependent memory traces during sleep stabilises and redistributes them to long-term neocortical sites (Born & Wilhelm, 2012; Squire et al., 2015). Low cholinergic tone, an absence of interfering sensory input, and highly synchronised thalamocortical activity during NREM stages 2 and 3 (N2/N3) provide a favourable environment for these processes (Gais & Born, 2004b, 2004a; Krugliakova et al., 2026). High-amplitude slow oscillations (SOs, ∼1 Hz) synchronise neocortical neuronal activity into hyperpolarised down- and depolarised up-states, temporally organising fast spindles (12-15 Hz) into their up-states (Mölle et al., 2011; Steriade, 2006). Spindles in turn cluster high-frequency hippocampal sharp wave-ripples (80-120 Hz) in their excitable troughs, which are considered to reflect memory trace reactivations (Staresina et al., 2015; Wilson & McNaughton, 1994). This hierarchical nesting of SOs, spindles and ripples provides a fine-tuned temporal framework for the hippocampal-neocortical communication and systems-level consolidation of memories (Diekelmann & Born, 2010; Staresina et al., 2023).

Within this cross-frequency coupling hierarchy, sleep spindles are considered to create transient windows for enhanced hippocampal-neocortical communication and synaptic plasticity in cortical circuits (Bergmann et al., 2012; Bergmann & Born, 2018). In mice, spindles have been found to coordinate the activity of distinct inhibitory interneuron populations, producing perisomatic inhibition while transiently disinhibiting the apical dendrites of pyramidal cells. The resulting synchronised, local Ca2+ influx fosters conditions conducive to synaptic plasticity (Niethard et al., 2018; Peyrache & Seibt, 2020; Seibt et al., 2017). In humans, transcranial magnetic stimulation (TMS) studies complement these cellular findings. Single-(spTMS) and paired-pulse TMS (ppTMS) over the primary motor cortex, combined with electromyography (EMG) recordings of contralateral hand muscles to quantify modulations of the motor-evoked potential (MEP) amplitude, provides a readout of corticospinal excitability (CSE) and its modulation by intracortical circuits, for example via GABA-A receptor-mediated short-interval intracortical inhibition (SICI) and glutamatergic intracortical facilitation (ICF) (Kujirai et al., 1993; Ziemann et al., 1996). CSE decreased with sleep onset and with deepening NREM sleep, while remaining stable during REM sleep (Avesani et al., 2008; Grosse et al., 2002; Manganotti et al., 2004; Salih et al., 2005). Accordingly, SICI increased from wakefulness to N3 sleep, whereas ICF remained unchanged but disappeared entirely during REM sleep (Avesani et al., 2008; Salih et al., 2005). Findings for lighter NREM sleep are inconsistent, potentially owed to low trial numbers, small samples, and stimulation-induced arousals in N1, but also to transient SO- and spindle-related changes in CSE and their difference in density across N2 and N3 sleep (Krugliakova et al., 2026).

Real-time EEG-triggered TMS now allows these transient changes to be examined at specific SO and spindle phases (Bergmann, 2018; Wischnewski et al., 2024). CSE was found to be higher during SO up-than down-states, and recently also to be suppressed during spindles relative to both post-spindle refractory periods and spindle-free N2/3 sleep (Bergmann et al., 2012; Hassan et al., 2025). Notably, this spindle-related suppression was phase-specific, being confined to the falling phase. Given the distinct configuration of inhibitory interneuron activity during spindles (Niethard et al., 2018; Peyrache et al., 2011), this spindle-phase-specific suppression may reflect local inhibitory network dynamics measurable through SICI.

In the present study, we therefore used real-time EEG-triggered spTMS and ppTMS over the primary motor cortex to measure CSE and SICI at four sleep spindle phases (peak, falling, trough, rising), during spindle-free N2/N3 sleep, and during pre-sleep wakefulness. We hypothesised that isolated spindles would reduce CSE and increase SICI relative to spindle-free N2/N3 sleep, with the strongest effects during the falling phase (Hassan et al., 2025). We further expected reduced excitability and increased SICI during sleep than during wakefulness (Avesani et al., 2008; Grosse et al., 2002; Manganotti et al., 2004; Salih et al., 2005). Exploratory analyses examined spindle trials coinciding with SOs, which were excluded from the primary spindle-phase analyses given their less frequent occurrence. Since fast spindles preferentially nest in the excitable SO up-states, we expected these trials to show higher CSE and lower SICI than isolated spindles (Bergmann et al., 2012; Mölle et al., 2011).

## Methods

### Participants

Twenty (12 female, 8 male) right-handed (Edinburgh Handedness Laterality Quotient = 78.13 ± 14.49%), Oldfield 1980) healthy young adults (23.90 ± 3.29 years, range = [19 31]) completed this study after providing written informed consent. The study was initially approved by the Ethics Committee of the Medical Faculty of the University of Tübingen (Ethics Votes No.: 349/2018BO2 and 810/2021BO2) and subsequently by the local Ethics Committee of the Rhineland-Palatinate Chamber of Physicians (Vote No. 2019-14679). Participants received moderate monetary compensation for their participation. We explicitly recruited participants who considered themselves to be good sleepers, and able to sleep in supine position without substantial movement or frequent arousals for 2-3 h in the nighttime, which was needed to enable stable positioning of the TMS-coil on target during sleep. Participants were recruited from the local Rhein-Main Universities student population (level of education 4.14 ± 1.67 ISCED 2011) based on the following inclusion criteria: participants had to be 18-45 years old, right-handed, physically and mentally healthy, have no history of neurological, psychiatric or sleep-related disorders, have no diagnosed or self-perceived attention or memory deficits, not take any medications that might affect cognition, sleep or drowsiness, not have consumed legal drugs for at least three months, have no history of misuse of alcohol or substance abuse, and no contraindications to TMS or EEG. EEG contraindications related to skin diseases or sensitivities, whereas TMS contraindications were based on (Rossi et al., 2021), including: not being of age, having experience or first-degree family history of epilepsy or seizures, history of head surgery, traumatic brain or spine injury, tinnitus, stroke or syncope, neurological or psychiatric diagnoses, magnetic objects on or inside the head, implants in the body, and no current pregnancy.

Eligibility was confirmed during a physician-led intake video screening, after which eligible participants attended an adaptation session before any experimental sessions. At the start of each session, participants completed another daily TMS safety questionnaire reviewed by the experimenter. This documented recent participation in non-invasive brain stimulation studies, the timing and quantity of consumed alcohol, caffeine, recreational drugs and medication use, headache that day, and sleep duration the previous night.

Participants were required to maintain a regular sleep-wake schedule for four weeks before participation and between sessions, precluding night-shift work and facilitating timely sleep onset after lights-out at ∼22:30. Across the two days preceding each session, participants completed a sleep diary, which included questions about sleep quality and duration, and alcohol and caffeine consumption. Participants were instructed to maintain their usual sleep-wake and consumption habits, except during a final 24 h abstinence period. During these 24 h, participants abstained from caffeine, alcohol, and any medications not approved by the study physician and avoided strenuous exercise, napping, and sleeping past 09:00. These measures promoted adequate sleep pressure, ensured substance-free sleep, and minimised the likelihood of compensatory recovery sleep at the laboratory.

### Experimental Design

This study employed a within-subjects design, with spTMS and ppTMS (SICI) repeatedly applied during N2/N3 sleep, each protocol time-locked to four sleep spindle phase-angles (peak (0°), trough (180°), rising (270°), and falling (90°) flank), as well as a spindle-free baseline condition 2 s post-spindle. The 10 conditions (2 TMS trial types x 5 target states) were presented in pseudorandomised order in multiple blocks of 60 trials (6 trials per condition per block) during each night. Since Hassan and colleagues (2024) had detected no difference in CSE between the immediate spindle refractory period, defined as 1.5 s post spindle-centre, and baseline periods of spindle- and SO-free N2/N3 sleep, we combined the two into one spindle-free baseline condition, for which we administered TMS 2 s post-spindle and post-hoc excluded any trials with new spindle-activity visible immediately preceding the timepoint of stimulation. Additionally, pre-sleep wake measurements of 40 trials per TMS protocol were obtained with protocol order counterbalanced across participants.

Participants attended one adaptation and 1-3 experimental sessions (1.80 ± 0.62) in the sleep laboratory at the Neuroimaging Center (NIC) of the Johannes Gutenberg University Medical Center Mainz, since data needed to be pooled across sessions to collect a minimum of 8 blocks (i.e., 48 trials per condition), aiming for 10-12 blocks, per participant (i.e., 60-72 trials per condition). Sessions were at least 5 days apart for sleep quality purposes. We intentionally chose to sample in blocks of 60 trials to account for gradual shifts in excitability and SICI across the night and, if necessary, recalibrate resting motor thresholds (RMT) between blocks (see Methods on TMS).

Sessions started around 20:00 and mostly ended between 01:30-02:30, with nocturnal naps lasting approximately 3 h, following a sleep onset latency of ∼30-minutes, and with TMS applied for the remaining 2.5 h. A session ended prematurely if participants woke and were unable to fall back asleep or wanted to leave.

### EMG, EEG and PSG

Surface EMG was recorded from three muscles of the left hand (abductor pollicis brevis (APB), first dorsal interosseous (FDI) and abductor digiti minimi (ADM)) using disposable sticky hydrogel electrodes (30 mm x 24 mm diameter) arranged in tendon-belly montages.

EEG was recorded from 64 TMS-compatible sintered Ag/AgCl ring electrodes (Multitrodes-TMS, EasyCap) embedded in a textile cap and arranged according to the 10-20 system (Fp1, Fpz, Fp2, AF7, AF3, AFz, AF4, AF8, F9, F7, F5, F3, F1, Fz, F2, F4, F6, F8, F10, FT9, FT7, FC5, FC3, FC1, FC2, FC4, FC6, FT8, FT10, T7, C5, C3, C1, Cz, C2, C4, C6, T8, TP7, CP5, CP3, CP1, CPz, CP2, CP4, CP6, TP8, P7, P5, P3, P1, Pz, P2, P4, P6, P8, PO7, PO3, PO4, PO8, O1, O2) and mastoid channels M1 and M2, with FCz as the reference, POz as the ground, and impedances kept under 5 kOhm.

For polysomnography (PSG), horizontal (HEOG) and vertical (VEOG) electrooculograms, as well as muscle tone from the chin (EMG) were recorded via additional bipolar channels and the same hydrogel EMG electrodes.

All signals were digitized in DC mode at a sampling rate of 5 kHz and with a 1250 Hz anti-aliasing low-pass filter via two 7.2V battery-powered 40 channel 24-bit amplifiers (NeurOne Tesla with Digital-Out Option, Bittium, Finland; NeurOne acquisition software v1.5.0).

### TMS

TMS was applied via an actively cooled butterfly coil with an outer diameter of 172x92 mm (MagVenture Cool-B65), and a stimulator capable of producing paired-pulse TMS (MagVenture MagPro X100 with MagOption). Participants lay in supine position for both wake and sleep measurements with the TMS coil placed tangentially over the right primary motor cortex. To ensure the coil handle was accessible from above (towards the participant’s forehead) and not digging into the mattress, the coil orientation, and accordingly the current direction set on the stimulator, was reversed by 180° (**Figure 1**), so that the induced current direction of the second half-wave of each biphasic pulse flowed in a posterolateral-anteromedial direction in the brain tissue.

**Figure 1.**
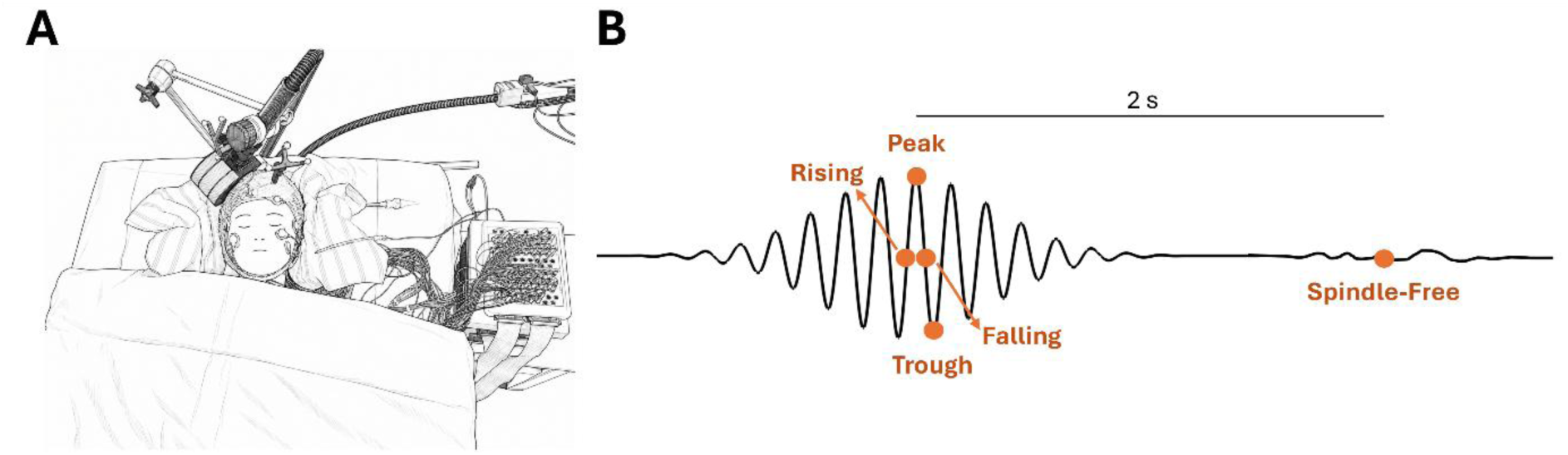
Real-time EEG-triggered TMS experimental setup and spindle-phase conditions. **(A)** Experimental setup of real-time EEG-triggered TMS in the sleep laboratory. The participant wears an EEG cap and lies in supine position with their head resting on a stabilising vacuum pillow. The TMS-coil is navigated onto the target (right primary motor cortex) and fixated using a coil holder. **(B)** The five stimulation conditions targeted with real-time EEG-triggered TMS during sleep include four spindle phases (rising, peak, falling trough) and a spindle-free N2/N3 stimulation condition occurring 2 s after a detected spindle. All conditions include only SO-free trials, identified offline.

Montreal Neurological Institute (MNI) template-based frameless stereotactic neuronavigation was used to track the coil and guide its positioning (Localite TMS Navigator v4.0.0, Germany; NDI Polaris Vega ST optical tracker). Participant’s supine head position was stabilised with a vacuum pillow (Vacuform) and the TMS coil was held in place via a mechanical arm attached to the bed (MagVenture Super Flex Arm) (**Figure 1**). During wake pre-measurements the tolerance for coil movement was < 1.5mm until the protocol was paused and the coil re-adjusted on target. In case participants moved off target during sleep by more than 1.5 mm or 1.5° and MEP amplitudes, which were continuously monitored, were visibly reduced (conditional tolerance zone), or latest if deviations exceeded 3 mm or 3°, stimulation was paused and an experimenter quietly entered the sleep cabin in semi-darkness to manually reposition the TMS coil without or only minimally disturbing the participant’s sleep.

We located the right individual motor hotspot as the coil position producing high-amplitude MEPs most consistently in any one of the three contralateral hand muscle EMG channels. This EMG channel was also selected as the target muscle for all subsequent TMS-EMG recordings. Starting with the coil placed above the right motor cortex tangentially to the scalp on electrode C4 with the induced current direction at a 90° angle to the precentral sulcus and a stimulation intensity of 40% maximum stimulator output (MSO), a repetitive systematic grid-search of the hand knob area was conducted, whereby the stimulation intensity of single pulses delivered at a jittered ISI of 3-4 s was increased in 5% steps whilst monitoring the EMG for MEPs. The coil position at the individual hotspot was saved and navigated back to for all subsequent stimulation protocols. The individual RMT was determined using an adaptive staircasing procedure implemented in the BEST-Toolbox (Hassan et al., 2022a). 40 spTMS pulses were delivered to the motor hotspot at a jittered ISI of 3-4 s, starting at the intensity that had elicited MEPs during the hotspot search, adapting it based on a >50 µV peak-to-peak MEP amplitude threshold. Participants were instructed to relax their left hand on top of a second vacuum cushion, and researchers monitored the chosen EMG-channel for precontraction. The RMT was defined as the intensity at which MEPs crossed this threshold during the last 5 of 10 applied pulses.

To measure CSE in the right primary motor cortex during pre-sleep wakefulness and N2/N3 sleep across target states, we applied spTMS at 120% of the individual RMT and measured peak-to-peak amplitudes in the EMG of the contralateral hand muscle (APB, FDI or ADM) individually selected during the hotspot search. Likewise, to quantify SICI, we applied paired pulse TMS to the right primary motor cortex with the conditioning stimulus (CS) at 80% RMT preceding the 120% wake-RMT test stimulus (TS) by 2 ms, and measured the percent reduction in MEP amplitude compared to the unconditioned spTMS (TS) MEPs (Di Lazzaro & Ziemann, 2013; Kujirai et al., 1993; Ziemann, 2004; Ziemann et al., 1996). During wake measures, the ITI was 5-6 s to prevent carry-over and expectancy effects. During sleep, the minimum inter-trial interval (ITI) was set to 6-7 s to prevent stimulation during any TMS-induced EEG activity and arousals. Ultimately, the ITIs depended on the natural individual occurrence of spindles.

We started stimulation during sleep using intensities calculated relative to participants’ pre-sleep wake RMT and closely monitored MEPs throughout each session. In sessions during which MEP amplitudes were so strongly suppressed during sleep, or gradually shifted across the night, that MEPs were no longer reliably measurable (i.e., no longer consistently exceeding 50 µV), especially during ppTMS trials (Avesani et al., 2008; Bergmann et al., 2019; Hassan et al., 2025; Salih et al., 2005), a short break was taken between blocks to reassess the RMT and adjust stimulation intensities for subsequent blocks. For efficiency, a quick manual thresholding procedure was run with single pulses applied at an ISI of ∼4-6 s starting at RMT + 3% MSO until the intensity was found at which 5/10 consecutive MEPs exceeded the 50 µV threshold.

All measurements were performed within a soundproof and electromagnetically shielded sleep cabin (Desone, Germany) equipped with a CO_2_ alarm and an infrared night-vision camera streaming to the control room, which participants were aware was for their safety. Participants lay in supine position on a comfortable mattress on a wooden bed with the EEG jackbox and amplifiers placed out of reach on a wooden nightstand. They were given TMS-compatible foam earplugs through which continuous active sound masking (TAAC-toolbox) was delivered to attenuate auditory co-stimulation and arousals induced by TMS-click sounds during sleep. The sound intensity was calibrated according to the participant’s preferences, ensuring that it could attenuate sound without preventing participants from falling asleep.

### Real-Time EEG-TMS

During experimental nights, individual phases (peak, trough, rising, and falling flank) of centroparietal sleep spindles in the right motor cortex were detected and targeted with TMS in real-time, as previously described in detail (Hassan et al., 2025; Hassan et al., 2022b). 64-channel EEG was streamed from the EEG recording system to the Brain Oscillation State Sensor (BOSS) device (bossdevice, sych2brain, Germany), a real-time PC for online signal processing. Spindles were detected via the Real-Time Spindle Detector (RTSD) algorithm (Hassan et al., 2022b) and their oscillatory phases via the *phastimate* algorithm (Zrenner et al., 2020), which were controlled using the Matlab software (2017b) based *Brain Electrophysiological recording and STimulation* (BEST)-Toolbox (Hassan, Pillen, et al., 2022). The RTSD algorithm was previously validated in the same laboratory as this study to accurately target spindle-phases consistently across participants, with a mean circular SD of 51° for all phases, resulting in ∼10 ms variations for an averaged 13 Hz spindle (Hassan et al., 2025).

A bipolar montage was chosen over a Hjorth montage to achieve higher signal-to-noise ratio given the synchronised nature of spindles across large areas of the cortex, like in a previous study on spindle-phase EEG (Hassan et al., 2025). From this channel, four signals with personalised thresholds were derived, of which 3 had to be surpassed for at least 250 ms to trigger a stimulation (for details see: Hassan et al., 2025). Therefore, spindles of ∼0.5-2 s duration were predominantly detected just after their centre, i.e. during their waning phase. We personalised the algorithm with adaptation night-derived individual mean root mean square (RMS) spindle power and peak spindle frequency, obtained offline using the Yet Another Spindle Algorithm (YASA) package run in Python software (Vallat & Walker, 2021) (see Analyses). Once detected in real-time, a spindle’s oscillatory phase was estimated on the fly using the *phastimate* algorithm (Zrenner et al., 2020), whereby the signal was autoregressively forward predicted (Yule-Walker 15^th^ order) for 65 ms to account for the hardware delay between event detection and TMS pulse delivery, and enable precise time-locking of the stimulus to the phase of interest.

### Procedure

#### Adaptation night

Upon arrival, participants were familiarised with the sleep cabins and study procedure before providing written informed consent. They then completed a demographic questionnaire and the Edinburgh Handedness Inventory (Oldfield, 1971). Participants unfamiliar with TMS received a brief demonstration to address any concerns. Otherwise, the adaptation night followed the same procedure as the experimental nights, except that no stimulation was delivered during sleep. To mimic the sensory conditions of experimental sessions, the TMS-coil was positioned on target, and noise masking was provided, but the stimulator remained off. After lights-out, participants had 2-3 h to fall asleep and transition into a second sleep cycle.

The adaptation night served three purposes. Firstly, it familiarised participants with the sleeping environment to reduce first-night effects and confirmed their ability to sleep well in supine position and without substantial head movement. A trained sleep stager continuously monitoring the PSG and TMS-coil position, and automated sleep staging was performed offline using YASA. Sleep onset latency, total sleep time, percentage wake after sleep onset, and time in N2/N3 sleep were assessed to confirm adequate sleep. Secondly, the adaptation night provided individual spindle parameters for personalised real-time detection during experimental nights. Using YASA’s automated spindle detection function *yasa.spindles_detect*() we extracted the mean individual peak spindle frequency and RMS deviation of spindles of 10-15 Hz, preferably fast spindles, detected at C4 during N2/N3 sleep identified via *yasa.SleepStaging*(). And thirdly, the session confirmed that the maximum planned stimulation intensity of 120% RMT would not exceed the MSO.

#### Experimental nights

Before each visit, sleep diary entries were reviewed to confirm adherence to study requirements. Participants arrived at the lab ∼20:00, completed the daily TMS-safety screening and prepared for bed. We applied the EEG, PSG, and hand EMG electrodes after disinfecting and lightly abrading the underlying skin, and checked signal quality. The EEG-cap was wrapped in cellophane and secured with a hairnet to prevent EEG-gel leaking onto the coil and pillow and bridging adjacent electrodes. Around 21:00, participants lay down in supine position for neuronavigation co-registration, and determination of the individual right motor hotspot and RMT.

Pre-sleep spTMS and ppTMS measurements were conducted around 22:00-22:30 while participants remained fully relaxed and fixated on the ceiling, after a short interactive break to maintain wakefulness. If participants’ eyes began to close or occipital alpha activity emerged, measurements were paused, and they were reminded to stay alert and offered water. Following a mandatory bathroom break and calibration of the masking sound intensity, the head position was stabilised with a vacuum pillow, and the TMS-coil held by the mechanical arm was repositioned on target. Participants were handed a bell to contact the experimenters before lights-out and closure of the cabin door ∼22:30.

After the first 5 min of stable N2 sleep, two AASM-trained researchers visually assed the accuracy of the real-time spindle detection, personalised with adaptation night-derived parameters. Fourty spTMS trials at 0% MSO were run to inspect the real-time placement of stimulation markers in the EEG/PSG. To control the algorithm’s detection sensitivity as advised, the RMS threshold was reduced if too many spindles were missed, or raised if false positives accumulated (Hassan et al., 2025; Hassan et al., 2022b).

Next, the stimulation intensity was gradually ramped up to minimise awakening. SpTMS was manually triggered at intervals of at least 6s, starting at 20% MSO and increasing in steps of 3-5% until the individually required maximal target intensity (120% RMT) was reached. Ramping was paused following signs of arousal or sleep stage transitions, and resumed when N2 sleep had re-stabilised. Depending on the participant, this took between one and several tens of minutes, sometimes requiring multiple attempts. Usually around 23:30 (i.e. 60min after lights-out), the main real-time spindle-phase triggered spTMS and ppTMS protocol began and continued for as many blocks as possible over ∼2-3 h of sleep.

Throughout the nocturnal nap, two AASM-trained researchers located in the control room alternated continuous monitoring of the PSG for signs of stage transitions, arousals and movement, the EMG for amplitude drift in the MEPs, and neuronavigation for coil displacement. Stimulation was immediately paused when necessary to keep participants asleep.

At the end of the nocturnal nap, usually ∼02:00-03:00, participants were woken up. All electrodes were removed and participants either left the laboratory or continued sleeping without the equipment.

### Analyses

#### Dependent variables

To assess CSE, the average peak-to-peak MEP amplitude of the selected contralateral EMG channel was calculated for each of the five phase conditions following spTMS applied to the primary motor cortex. For ppTMS, SICI was expressed as %change in amplitude of the conditioned (CS+TS) from the unconditioned (TS) MEP peak-to-peak amplitude (% change = (SICI_ratio - 1) * 100), with more negative values reflecting stronger inhibition. EMG and EEG data was analysed in Matlab (v2023b) using the FieldTrip toolbox (v20240129) (Oostenveld et al., 2011), PSG data was sleep staged in Python (v3.13.3, Spyder v6.0.5) using the YASA package (v0.6.5) (Vallat & Walker, 2021), and statistical tests were conducted in R (v4.5.1, R-Studio v2025.05.1+513).

#### MEP amplitudes

EMG data was extracted from the session-specific muscle channel into epochs of -0.5 to 1.01 s around each trial’s TMS (TS) pulse, and baseline-corrected to a pre-TMS period -100 to -5 ms to rectify the DC offset. To extract the peak-to-peak MEP amplitude, a DFT notch filter for line noise of 50, 100 and 150 Hz was applied to a post-TMS segment of 0.01 to 1.01 s, and the sum of the absolute minimum and maximum value within a 20 to 50 ms post-TMS window was computed.

#### SICI ratios and normalised MEPs

To account for stimulation-intensity changes across blocks and slow drifts in baseline CSE across sleep stages and the progressing nap, we normalised MEPs and calculated SICI ratios within shorter data segments consistent with previous real-time TMS-EEG studies (Bergmann et al., 2012; Hassan et al., 2025; Thies et al., 2018; Zrenner et al., 2018). Segments were designed to provide stable condition estimates while remaining sensitive to temporal drift in CSE. Each of the 60-trial blocks contained six trials per condition, of which ∼20% were expected to be excluded from the primary isolated spindle analyses because of coincident SOs. We therefore combined multiple blocks acquired at the same intensity into more reliable and representative segments. Where possible, each segment comprised four blocks, containing 24 trials per condition before trial exclusion, consistent with previous studies, where blocks had contained ∼15-20 trials per condition. Segment size was kept consistent within each session to maintain comparable variance, for example, a session with 6 blocks was divided into two three-block segments rather than segments of four and two blocks. Segments were only shorter if fewer blocks at the same intensity were available, typically before any intensity adjustments early in the session.

The geometric mean was used to estimate the central tendency of each condition within each segment, because MEP amplitudes were positively skewed and log-normally distributed. The geometric mean corresponds to exponentiating, i.e., back-transforming, the arithmetic mean of log-transformed MEP amplitudes, whilst retaining the more interpretable original µV scale. Hereafter, the “mean” refers to the arithmetic mean, unless the geometric mean is explicitly specified.

Within each segment, normalised MEPs were calculated as the percentage change of each condition’s geometric mean relative to the mean across all spTMS conditions’ geometric means. SICI was calculated as a ratio of the relevant geometric condition means within each segment:

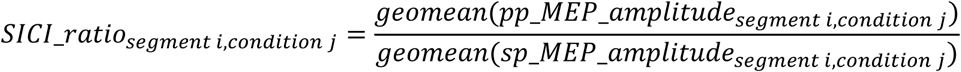

To compute the session means, each segment was weighted by the number of blocks it had been constructed from, and hence the period of the recording it represented. If data was pooled across multiple sessions, different recording lengths were equally accounted for in the condition means of each participant.

#### Trial exclusion

Trials contaminated by EMG precontraction, movement- or arousal-related EEG artifacts, or spindle activity during nominally spindle-free post-spindle trials were excluded. To isolate spindle-phase effects from the known modulation of excitability by SO phase (Bergmann et al., 2012), trials with SOs present at stimulation were also excluded from the primary analyses.

To detect EMG precontraction, data from -500 to -1 ms of the initial EMG epoch was demeaned, detrended, bandpass-filtered at 10-500 Hz, and DFT notch-filtered at 50, 100, and 150 Hz. Precontraction was defined as the RMS of 200 ms pre-TMS signal exceeding a session-relative threshold of mean RMS + 2 SD.

For EEG-based exclusions, pre-TMS EEG-activity from the conjunction of a local C4-area montage (C2, C4, C6, FC4, CP4), left-mastoid (M1), and selected lateral channels sensitive to cranial muscle and movement artifacts (Fp1, Fp2, F7, F8, T7, T8, P7, P8, O1, O2), was epoched from -1 to -0.001 s and demeaned. If M1 was extremely noisy, it was replaced by a clean neighbouring electrode (e.g. TP7). Noisy EEG trials were excluded due to the ambivalence of spindles detected from a contaminated signal. To identify high-frequency contamination, the lateral channel epochs were zero padded by 0.2 s on each side and bandpass-filtered at 40-1000 Hz. The Hilbert transform extracted the signal envelope, and trials with an envelope z-score exceeding 15 were excluded.

Post-spindle “spindle-free” condition trials were manually inspected by an expert AASM-trained sleep stager (Iber et al., 2007). For inspection, -3 to 2.5 s EEG epochs around each TMS-pulse were extracted from the standard PSG channels (F4, C4, P4, O2 and M1) and the C4-area montage, re-referenced to the contralateral mastoid (M1) like during real-time detection, and demeaned. Trials with spindle activity unambiguously visible in C4 and surrounding channels at or within 0.5 s of stimulation, typically reflecting new spindle activity ∼2 s after the detected spindle, were excluded. Like for SOs, automated detection using YASA’s *yasa.spindle_detect*() was unreliable because the high-amplitude TMS-pulse artifact contaminated the signal from the middle of the targeted spindle onwards.

To detect SO-presence at stimulation, the pre-TMS EEG epochs of C4-area channels were 40 Hz low-pass filtered, re-referenced to linked mastoids, and demeaned. The minimum signal amplitude was extracted from each channel to identify negative SO peaks. Candidate SOs exceeded a session-relative threshold of the mean absolute negative peak + 0.5 SD, capped at 80 µV to accommodate sessions with high N3 sleep proportions. Trials in which SO-candidates were detected in at least three of the C4-area channels were classified as containing an SO. This prevented false detections caused by respiration-related, high-amplitude slow artifacts in one or two channels in contact with the TMS-coil. In case two or more C4-area channels were unavailable, frontal channels F2, F4, and F6 were included, and agreement across at least two channels was required.

This procedure was developed and validated across participants by an expert sleep stager based on hits, misses, false positives, false negatives. It outperformed automated SO detection using YASA’s *yasa.sw_detect()* run on EEG data with TMS-contaminated periods removed and interpolated. Standard morphological, duration, fixed amplitude (> ∼75 µV), and zero-crossing criteria were unsuitable because the TMS-pulse truncated the SO waveform at variable phases. Whilst spindles usually nest in the SO up-state, we did not want to restrict our detection to fully formed SOs considering that stimulation could occur at any SO-phase in the spindle-free condition. The short pre-TMS window ensured identification of SOs ongoing at stimulation, while the local C4 montage ensured relevance to the motor cortex.

#### Data for SO-presence analyses

Trials with SOs were excluded from all analyses except for the specific tests of SO-nested vs. isolated spindles. Data from all clean trials containing SOs was split into a spindle-average condition with SO-nested spindle trials and a spindle-free N2/N3 sleep condition containing isolated SOs. Since SOs were expected to arise spontaneously and unevenly across segments and conditions, with the probability of nested SOs increasing during periods of N3 sleep, condition geometric means were computed across all raw MEP amplitudes of a session instead of being normalised within each segment. SICI ratios were also calculated at the session level.

#### Data for wake vs. sleep analyses

Peak-to-peak MEP amplitudes from pre-sleep wakefulness measurements were extracted from segments of EMG data spanning -50 to 150 ms around each TMS pulse, saved in the BEST-Toolbox output of each session, following the same procedure as for sleep MEPs. Trials in which amplitudes in the 50 ms preceding each TS pulse exceeded a relative threshold of 25 µV above the average of the ten smallest pre-TMS peak-to-peak amplitudes of that protocol and participant were deemed to be noisy or include precontraction and were excluded from the respective 40-trial average. The geometric means were calculated for each wake protocol and session, and SICI was calculated at the session level as their ratio.

Regarding the sleep data, only blocks of data acquired at the same stimulation intensities as during pre-sleep wakefulness were included in this SO-free spindle-average condition to ensure fair comparison to wake data. This reduced both sample size and trial numbers, as intensity increases had been required in 12/20 participants, usually towards the beginning of the sleep recording and sometimes even before the first block of data was collected. Sessions with a minimum of 10 clean spindle-condition trials per TMS protocol were included for the reduced sleep spindle dataset, and sessions contributed equally to the geometric subject means. To complete the picture, the same procedure was performed to obtain a spindle-free dataset acquired at preceding wakefulness stimulation intensities, despite lower trial numbers.

#### Sleep staging

Before automatically sleep staging polysomnography data using YASA (Vallat & Walker, 2021), TMS-induced artifacts in the EEG data had to be removed. The following procedure was confirmed to achieve high-quality automated sleep staging results in YASA compared to visual staging. PSG data from a common bilateral sleep staging montage was extracted (channels F3, F4, C3, C4, O1, O2, M1, M2, HEOG, VEOG, EMG), segments of [-20 +50] ms around each TMS trial were linearly interpolated based on 10 ms of neighbouring data from each side of the segment. The data was downsampled to 100 Hz, re-referenced to M1, bandpass filtered ([0.1 40] Hz) and automatically sleep staged including confidence estimates in Python via YASA’s *yasa.SleepStaging*() function, based on channel C4, or if noisy C2, an EOG and EMG channel (HEOG, chin-EMG).

### Statistics

The following statistical analyses were conducted separately for MEP amplitude and % SICI. A significance level of α = 0.05 was applied for all statistical tests, and effect sizes for all F-tests reflect partial eta-squared, and Cohen’s d for t-tests. Significance levels are indicated as * p < .05, ** p < .01, and *** p < .001. All measures of central tendency and variance reflect mean ± SD, and all error bars in figures represent the standard error of the mean (SEM), unless stated otherwise. Numbers are rounded to two significant digits.

To test whether MEP amplitude or SICI changed across spindle-phase, we ran linear mixed-effects models fitted with restricted maximum likelihood (REML) using R package lmerTest’s function *lmer()*, including spindle-phase condition (5 levels: peak, trough, falling, rising, spindle-free) as a fixed effect and a random intercept for participant to account for subject-specific variance across repeated measures: MEP_amplitude ∼ phase_condition + (1 | subject), and SICI ∼ phase_condition + (1 | subject), respectively. Normality of residuals was assessed with normal Q-Q plots, and homogeneity of variance by plotting residuals against fitted values. We conducted a Type III analysis of variance F-test on the model’s fixed effects using the function *anova()*, with denominator degrees of freedom estimated using Satterthwait’s method. Following a significant omnibus effect of spindle-phase condition, post-hoc pairwise comparisons were conducted on the model-estimated marginal means calculated using the emmeans package’s *emmeans()* function. P-values were adjusted for multiple comparisons via the Benjamini-Hochberg family-wise error correction, and Cohens’ d effect sizes were calculated using the emmeans package’s *eff_size()* function. Based on previous findings (Hassan et al., 2025), we hypothesised that MEP amplitudes would be significantly smaller during the falling phase condition than during all other conditions, accompanied by selectively more negative %change SICI values (i.e., stronger inhibition).

To test for an effect of spindle presence, we ran a one-sided paired t-test on the model’s estimated marginal means comparing the spindle-average to the spindle-free condition. The mean of four spindle-phase condition means (peak, trough, rising and falling flank) was taken to represent the derived spindle-average condition for each participant. In agreement with previous results, we hypothesised that spindle-presence would relatively reduce MEP amplitude (Hassan et al., 2025) and accordingly lead to more negative %change SICI values.

All previous analyses were conducted on isolated spindles only, as the percentage of SO-nested spindles was expected to be too low to test for reliable phase-effects. However, to test for an effect of SO-nesting on MEP amplitude and SICI, a two-sided paired t-test of isolated vs. SO-nested spindles was conducted on the derived spindle-average conditions. We hypothesised that amplitudes would be larger and inhibition smaller during SO-nested spindles, given the relative increase in MEP amplitude during the SO up-state, during which spindles are mostly nested, compared to down-state (Bergmann et al., 2012). For a complete picture, an equivalent two-sided t-test was conducted on the two spindle-free conditions, one where SOs were present, the other with neither SOs nor spindles present. Since trial numbers of spindle-free SO-nested trials were expected to be very low, this comparison should be considered as exploratory.

To test for the hypothesised reduction in MEP amplitude from wake to sleep and accompanying increase in inhibition (more negative % SICI values), as found in several previous studies (Avesani et al., 2008; Bertini et al., 2004; Grosse et al., 2002; Manganotti et al., 2004; Salih et al., 2005), a one-sided paired t-test compared the pre-sleep wakefulness to the spindle-average condition. To complete the picture an equivalent test in the same direction compared pre-sleep wakefulness to the spindle-free condition, although lower trial numbers were expected in these conditions.

## Results

### Sample

To reach the target sample size of 20 participants, 59 participants took part in an adaptation night, of which 38 returned for an experimental night, respectively of which 16 were excluded or dropped out due to poor sleep quality, the inability to remain asleep during ramping up to the required stimulation intensity, or their unavailability to return for further sessions to obtain sufficient trials. The final dataset was collected across 1 to 3 experimental sessions per participant (mean ± SD, 1.77 ± 0.53), with 8 to 17 blocks of data per participant (12.45 ± 2.058), and 30 to 82 clean SO-free MEPs per condition (54.91 ± 8.90) (**Table 1**). Trials contaminated by EMG-precontraction, noisy EEG, or spindle activity during post-spindle trials were rejected (2.83 ± 1.53%) and, to isolate spindle-phase related effects from the known modulation of excitability by SO-phase (Bergmann et al., 2012), trials with SO present at stimulation (21.31 ± 4.71%) were also excluded from the main analyses.

**Table 1.**
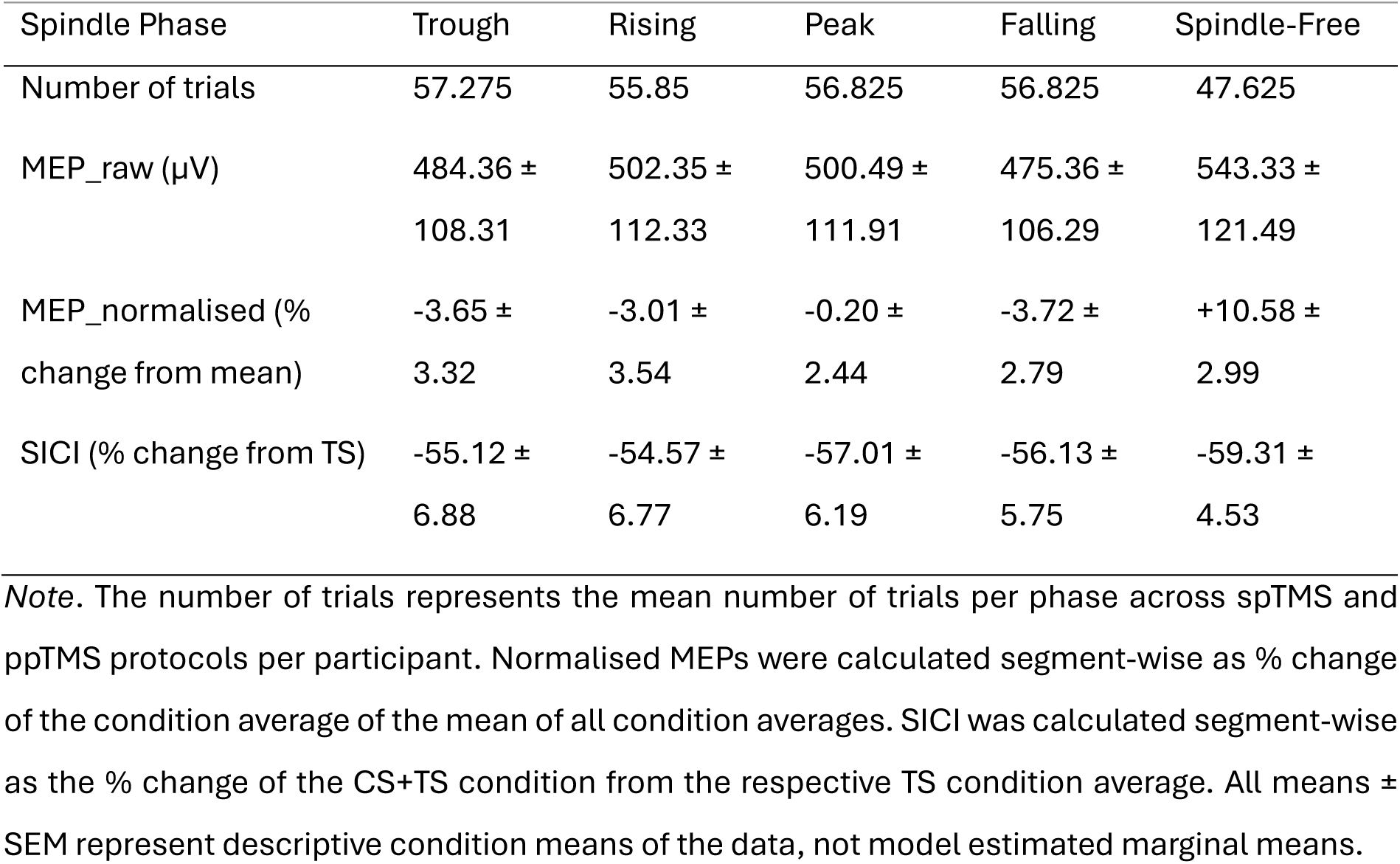
Trial counts, descriptive mean ± SEM of spTMS MEP amplitudes, SICI, and conditioned ppTMS MEP amplitudes, per spindle-phase condition.

To combat floor effects for SICI, resulting from strongly suppressed spTMS MEP amplitudes (TS), RMTs were recalibrated during sleep and stimulation intensities increased in 12 of 20 participants, who presented especially strong sleep-induced suppression of MEP amplitudes (spTMS mean < 50 µV; mean increase = 4.04 ± 2.55 % MSO per adjusted session). Consequently, stimulation intensities during sleep were slightly higher (80% and 120% respectively of individual RMTs of 61.04 ± 9.48 % MSO) than during preceding wakefulness (RMT = 58.76 ± 9.79 % MSO). The average individual spindle frequency was 12.37 ± 0.47 Hz. Descriptive condition averages for MEP amplitudes and SICI are summarised in **Table 1**.

### spTMS MEP amplitudes but not SICI modulated by sleep spindles

SpTMS MEP amplitudes were significantly modulated by spindle-phase condition (*F*(4,95) = 4.01, *p* = .0048, *η_p_*^2^ = 0.14; **Figure 2A**). The strongest suppression from the mean across all conditions (normalized MEP) occurred during the falling phase and the trough, followed by the rising phase, and with only little suppression during the peak, whereas only the spindle-free condition displayed a relatively larger normalized MEP amplitude (see **Table 1** for mean values). During all four within-spindle phase conditions, MEP amplitudes were significantly lower than during the spindle-free condition (*Trough*: MD = -14.24%, 95% CI = [-26.60 -1.87], *t*(95) = -3.31, *p_adj_* < .01, *d* = -1.047; *Rising*: MD = -13.59%, 95% CI = [-25.95 -1.23], *t*(95) = -3.16, *p_adj_* < .01, *d* = -1.00; *Peak*: MD = -10.78%, 95% CI = [-23.14 +1.58], *t*(95) = -2.51, *p_adj_* < .05, *d* = -0.79; *Falling*: MD = -14.30%, 95% CI = [-26.66 -1.94], *t*(95) = -3.32, *p_adj_* = < .01, *d* = -1.051). No significant differences in MEP amplitude were found between any of the four within-spindle phase conditions [all: *p_adj_* >.70].

**Figure 2.**
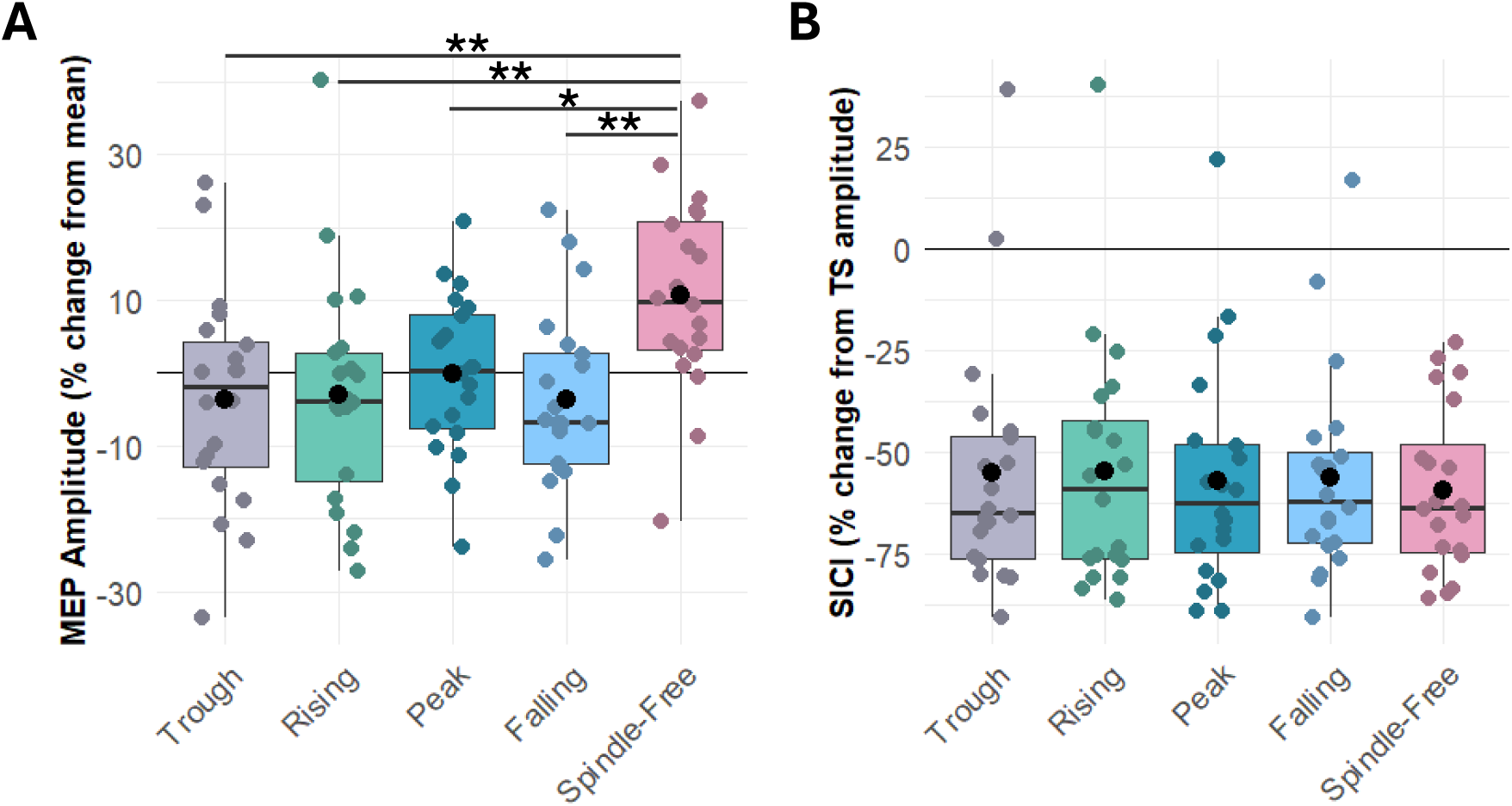
MEP amplitude and SICI as a function of spindle-phase. (***A***) Normalised MEPs are presented as % change in amplitude from the mean across all conditions within a normalisation block to account for drifts over time and adjustments in stimulation intensities. MEP amplitudes were modulated by spindle events, showing a significant reduction in all spindle-phase conditions compared to the spindle-free N2/N3 condition. While MEP amplitudes were numerically smallest for the falling phase condition, no significant difference was observed between spindle phases (*p_adj_* > 0.70). Significance of BH-corrected post-hoc comparisons is indicated as follows: * *p_adj_* < .05, ** *p_adj_* < .01, *** *p_adj_* < .001. **(*B*)** SICI is presented as % change in the CS+TS MEP amplitude from the unconditioned TS amplitude within a normalisation block. While SICI was clearly present during N2/N3 sleep across all spindle and spindle-free conditions, SICI was not significantly modulated across spindle-phase conditions (*p* = .80). Boxplots show the distribution of values for each condition, with the box spanning the interquartile range (IQR), the horizontal line indicating the median, and the black circle indicating the mean. Jittered coloured dots represent single-subject values. The horizontal reference line at y = 0 indicates no change in SICI.

The effects in MEP amplitude were not mirrored by SICI. While clear SICI was strongly expressed during all five spindle-phase conditions (*Trough*: M = -55.10%, 95% CI = [-67.60 -42.70]; *Rising*: M = -54.60%, 95% CI = [-67.00 -42.10]; *Peak*: -57.00%, 95% CI = [-68.60 -43.70]; *Falling*: M = -56.10%, 95% CI = [-68.60 -43.70]; *Spindle-Free*: M = -59.30%, 95% CI = [-71.8 -46.9]; all one-sided t-tests against 0: p_adj_ < .001), SICI was not significantly modulated as a function of spindle-phase [*F*(4, 76) = 0.41, *p* = .80, *η_p_*^2^ = 0.02] (**Figure 2B**).

Averaged across the four within-spindle phase conditions, spTMS MEP amplitudes were significantly suppressed during spindles compared to spindle-free N2/N3 sleep, consistent with our hypothesis (MD = -13.20, 95% CI = [-20.00 -6.47], *t*(95) = -3.89, *p* = < .001, *d* = -0.87) (**Figure 3A**). In contrast, SICI was not significantly different during spindles than spindle-free N2/N3 sleep (MD = 3.61%, 95% CI = [-2.88 +10.10], *t*(95) = 1.11, *p* = .27, *d* = 0.25) (**Figure 3B**).

**Figure 3.**
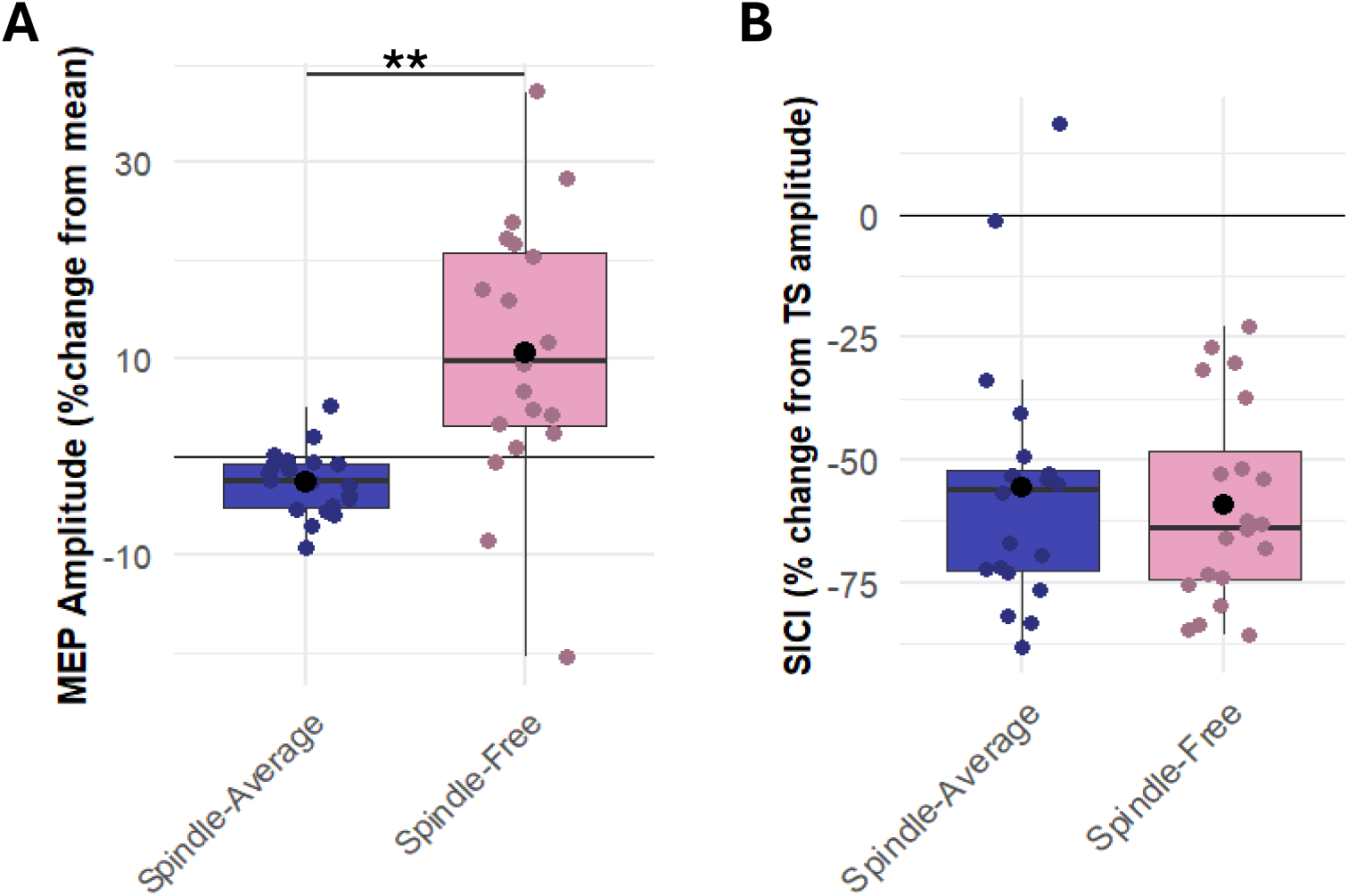
MEP amplitude and SICI during spindles and spindle-free N2/N3 sleep. (***A***) MEP amplitudes were significantly suppressed by spindle-presence compared to spindle-free N2/N3 sleep. **(*B*)** Whilst SICI was strongly expressed both during spindles and spindle-free N2/N3 sleep, it was not significantly affected by spindle-presence. Significance of BH-corrected post-hoc comparisons is indicated as follows: * *p_adj_* < .05, ** *p_adj_* < .01, *** *p_adj_* < .001. Boxplots show the distribution of values for each condition, with the box spanning the interquartile range (IQR), the horizontal line indicating the median, and the black circle indicating the mean. Jittered coloured dots represent single-subject values. The horizontal reference line at y = 0 indicates no change in SICI.

### SO-nesting of spindles increases MEP amplitude and reduces SICI

As hypothesised, MEP amplitudes were significantly larger when spindles were nested in SO than during isolated spindles (MD = 211.10 µV, 95% CI = [98.99 323.20], *t*(19) = 3.94, *p* = < .001, *d* = 0.30) (**Figure 4A**). The same pattern emerged in the absence of spindles, where MEP amplitudes were significantly larger during SO presence than during SO-free and spindle-free N2/N3 sleep (MD = 124.49 µV, 95% CI = [8.78 Inf], *t*(19) = 3.07, *p* = < .01, *d* = 0.19).

**Figure 4.**
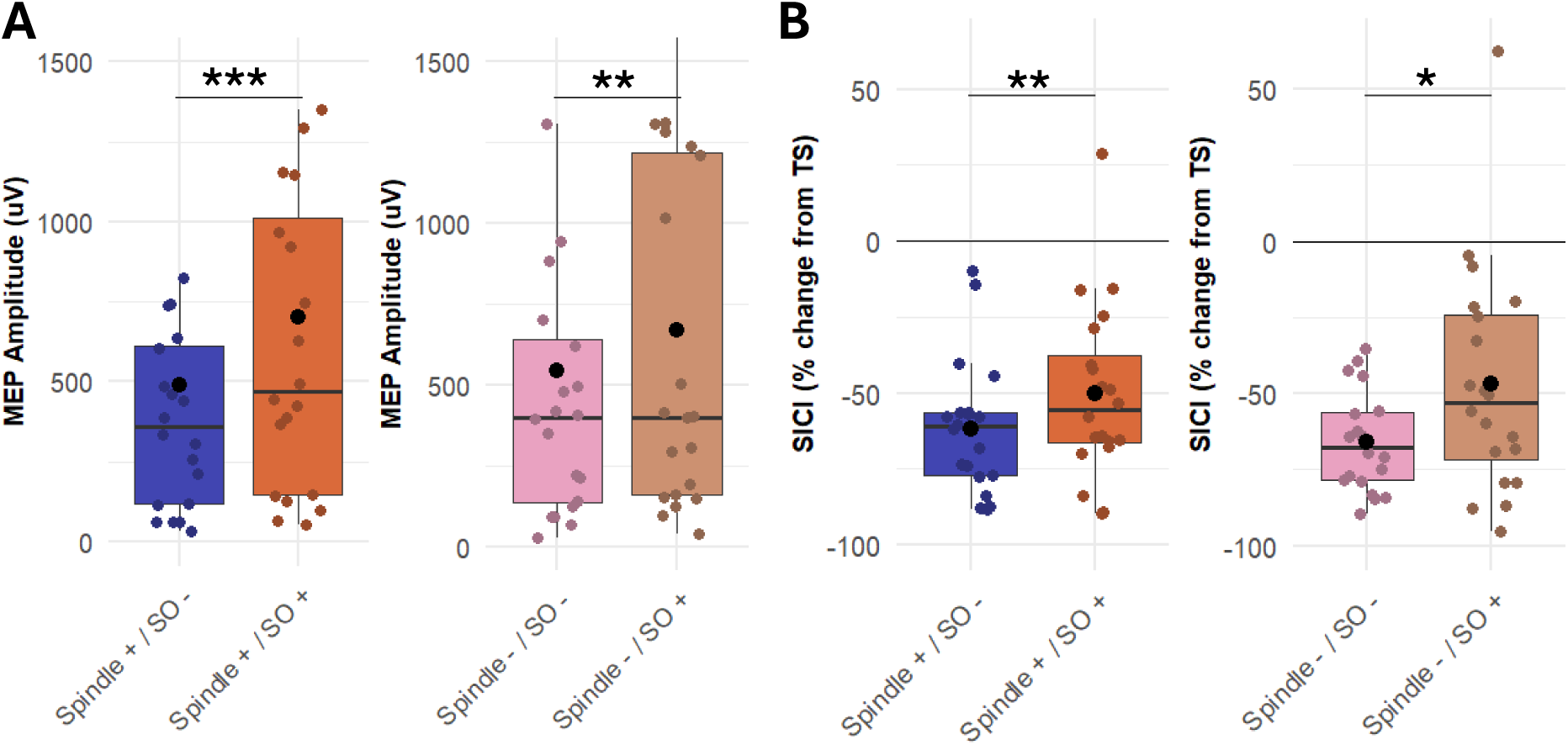
MEP amplitude and SICI across SOs and spindle presence. (***A***) SO presence, whether in the presence (+) or absence (-) of spindles, significantly increased MEP amplitudes. Note one outlying subject with amplitudes exceeding 2.5mV across all conditions was omitted from the axis to preserve detail for the remaining data. **(*B*)** SO presence, whether in the presence or absence of spindles, also significantly reduced the strength of SICI. Significance of BH-corrected post-hoc comparisons is indicated as follows: * *p_adj_* < .05, ** *p_adj_* < .01, *** *p_adj_* < .001. Boxplots show the distribution of values for each condition, with the box spanning the interquartile range (IQR), the horizontal line indicating the median, and the black circle indicating the mean. Jittered coloured dots represent single-subject values. The horizontal reference line at y = 0 indicates no change in SICI.

As hypothesised, this effect was mirrored by SICI (**Figure 4B**). When spindles were nested in SO, inhibition was reduced (less negative % SICI values) compared to isolated spindles (MD = +11.29 %, 95% CI = [+3.60 +19.00], *t*(19) = 3.07, *p* = < .01, *d* = 0.41). Similarly, in the absence of spindles, the strength of inhibition was reduced during SO presence compared to SO-free and spindle-free N2/N3 sleep (MD = +18.93 %, 95% CI = [+34.77 +3.08], *t*(19) = 2.50, *p* = < .05, *d* = 0.60). The two tests of SO presence in the absence of spindles should be understood as exploratory given their comparably lower trial numbers (trial numbers for each TMS protocol: spindle-free: 8.98 ± 4.54; spindle-average: 63.90 ± 20.55).

### From wake to N2/3 sleep (spindles) MEP amplitude is suppressed and SICI enhanced

To ensure an unbiased wake-sleep comparison, the following results are based on two separate reduced samples (spindle-average: n = 18, spindle-free: n = 14) as detailed in the methods, only including SO-free data acquired at the same stimulation intensities as during preceding wakefulness where available (number of trials per participant for each TMS protocol: spindle-average = 132.45 ± 78.80; spindle-free: 32.26 ± 11.73). As hypothesised, MEP amplitudes were strongly suppressed during sleep spindles (MD = -839.85 µV, 95% CI = [-Inf -571.66], *t*(17) = -5.45, *p* = < .001, *d* = -0.52) (**Figure 5A**), as well as during spindle-free N2/N3 sleep compared to preceding wakefulness (MD = -840.50 µV, 95% CI = [-Inf -422.76], *t*(13) = -3.56, *p* < .01, *d* = -0.63) (**Figure 5C**).

**Figure 5.**
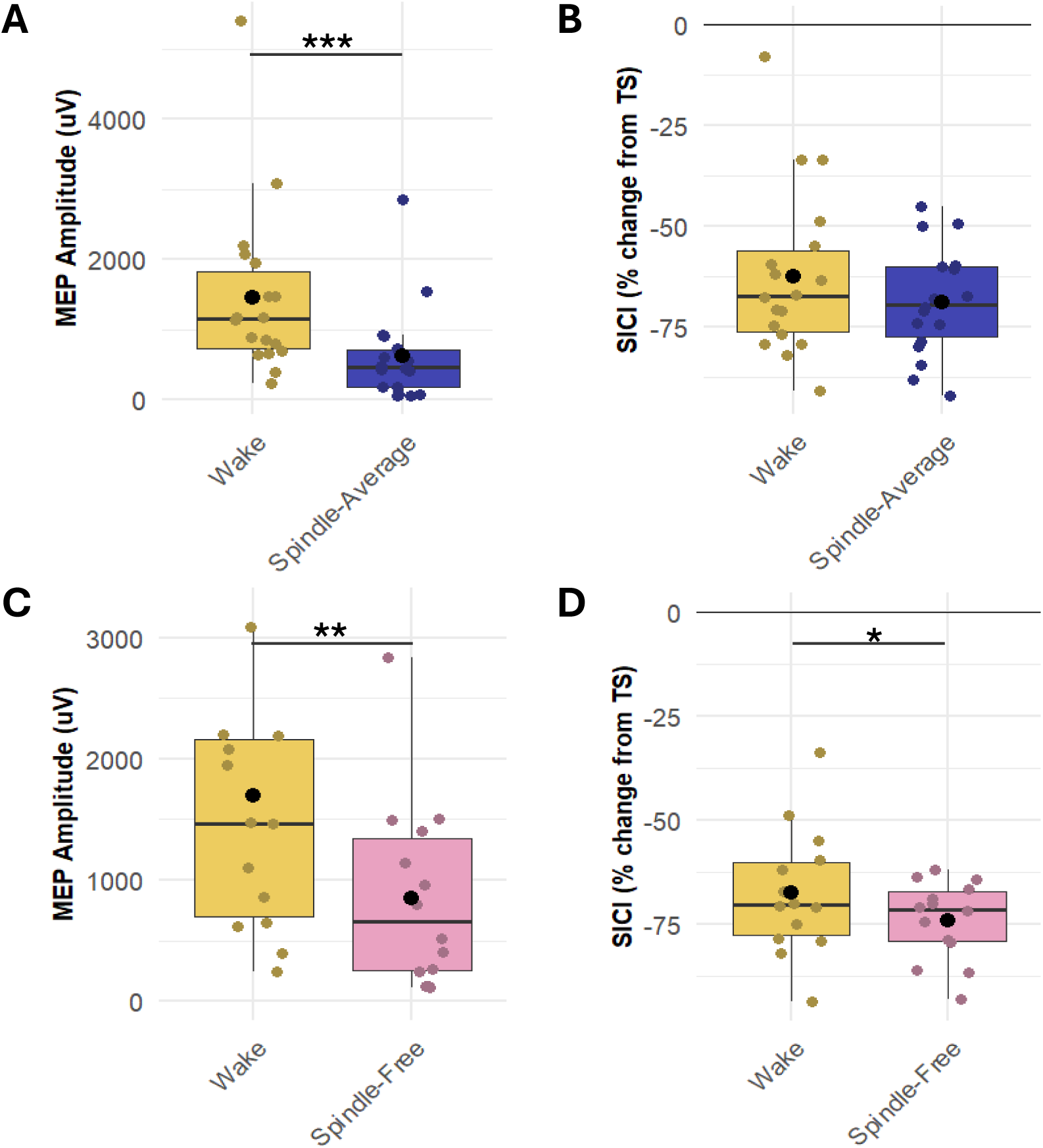
MEP amplitude and SICI from wakefulness to sleep (spindles). (***A-B***) During sleep spindles, MEP amplitudes were significantly suppressed relative to preceding wakefulness, whereas SICI was not significantly stronger (subsample n = 18). (***C-D***) During spindle-free N2/N3 sleep, MEP amplitudes were also significantly suppressed relative to preceding wakefulness, accompanied by a significant increase in SICI strength (subsample n = 14). Significance of BH-corrected post-hoc comparisons is indicated as follows: * *p_adj_* < .05, ** *p_adj_* < .01, *** *p_adj_* < .001. Boxplots show the distribution of values for each condition, with the box spanning the interquartile range (IQR), the horizontal line indicating the median, and the black circle indicating the mean. Jittered coloured dots represent single-subject values. The horizontal reference line at y = 0 indicates no change in SICI.

These results were accompanied by weaker changes in SICI. We found a non-significant trend towards an accompanying increase in the strength of SICI from pre-sleep wakefulness to sleep spindles (MD = -6.56% µV, 95% CI = [-14.60 +1.48], *t*(17) = -1.72, *p* = .052, *d* = -0.52) (**Figure 5B**), and a marginally significant increase in inhibition from wakefulness to spindle-free N2/N3 sleep (MD = -6.45% µV, 95% CI = [-14.30 -1.40], *t*(13) = -1.77, *p* < .05, *d* = -0.52) (**Figure 5D**).

## Discussion

This study characterised sleep spindle-related changes in corticospinal excitability (CSE) and GABA-A receptor-mediated inhibition in humans using real-time EEG spindle-phase-triggered TMS and obtained four main findings. Firstly, we replicated the reduction in CSE during spindles relative to a spindle-free N2/N3 sleep (Hassan et al., 2025). Contrary to the previously reported spindle falling phase-specific effect however, we found that reduction occurred across all spindle phases, although the numerically strongest effects were again exerted by the falling flank and trough. Secondly and contrary to our hypothesis, neither spindle presence nor phase significantly modulated SICI, suggesting that the spindle-related suppression of CSE was not accompanied by disproportionally high GABA-A receptor-mediated inhibition. Thirdly, CSE was strongly reduced from preceding wakefulness to both spindle-free epochs as well as spindles during N2/N3 sleep, whereas SICI only showed a trend towards a respective increase during sleep, which is in line with previous mixed findings on SICI. Finally, secondary analyses provided preliminary evidence that CSE was increased when SOs were present, both in the presence or absence of spindles, in accordance with the excitability increase previously observed during the SO up-state relative to the down-state (Bergmann et al., 2012), and that this effect was accompanied by a reduction in SICI during SOs. Together, these findings reveal distinct contributions of spindles, SOs, and the general vigilance state of sleep vs. wakefulness to CSE and GABA-A receptor-mediated inhibition.

### Reduced excitability during sleep spindles

In line with Hassan and colleagues (2025), we found that isolated sleep spindles reduced CSE relative to spindle-free N2/N3 sleep. In our data, this reduction was not driven solely by the falling phase, but by all spindle phases, although we also found the numerically largest reductions in excitability during the spindle falling phase, as well as the trough.

In absence of the previous study, one might have expected a trough-specific facilitation, given that this is the oscillatory phase in which spindles nest the hippocampal ripples, given that hippocampal ripples cluster in spindle troughs, potentially creating phase-specific windows for reactivation and plasticity (Dickey et al., 2021; Staresina et al., 2023). The trough, or early rising phase, was also shown to be the phase of highest excitability in the sensorimotor mu-alpha oscillation during wakefulness (Bergmann et al., 2019; Desideri et al., 2019; Schaworonkow et al., 2019; Stefanou et al., 2018; Zrenner et al., 2018, 2023). Whilst not significant, our data reflects this numerically, with the strongest suppression in CSE exhibited during trough, and falling phase. It is possible that the true most excitable phase lies in between falling phase and trough, and that by re-binning the phase conditions to fit tighter distributions around each phase angle (0°, 90°, 180°, 270°), differences between individual spindle-phase conditions could become more pronounced.

No significant evidence in favour of a phase-specific modulation of CSE does not exclude potentially phase-dependent cortical microcircuit dynamics in line with the literature, but that any effects proportionally offset each other and not change the overall excitability of the corticospinal system. Specifically, spindles are thought to phasically decrease the activity of somatostatin (SOM+) inhibitory interneurons, whilst increasing the activity of parvalbumin (PV+) inhibitory interneurons, inducing a local configuration of dendritic disinhibition and periosomatic inhibition of pyramidal cells. This local compartmentalisation of an influx of Ca2+ in the apical dendrites, rhythmically synchronised to temporally precise and reoccurring windows during each spindle cycle, is thought to create a particularly favourable environment for the induction of synaptic plasticity (Brécier et al., 2021; Dickey et al., 2021; Klinzing et al., 2019; Niethard et al., 2018; Peyrache & Seibt, 2020; Seibt et al., 2017). Since the modulation of different inhibitory interneuron populations is bidirectional, these cycle-to-cycle changes at the local dendritic level may not induce any net changes in excitability at the level of the corticospinal system. This might explain why our results show a sustained level of CSE across spindle phases.

The reduction in excitability that we found to be sustained across all spindle-phases relative to spindle-free sleep may also reflect a transient state of cortical disengagement associated with spindles. Thalamic gating proposes that spindle-related activity blocks or at least attenuates the relay of sensory input into the cortex, a process mediated by thalamocortical circuits and GABAergic activity in the thalamic reticular nucleus (Fernandez & Lüthi, 2020; Lüthi, 2014). Our findings might reflect this spindle-related modulation of cortical responsiveness, although evidence on auditory transmission to the cortex during spindles has been mixed (Dang-Vu et al., 2011; Jourde & Coffey, 2024).

It is also possible that this study failed to detect a phase-specific modulation due to any of the following methodological differences to the previous study (Hassan et al., 2025). (i) We detected spindles using the same hardware, software, real-time detection algorithm and montage, but in the opposite, non-dominant, hemisphere (C4-M1 vs C3-M2). Therefore, we also stimulated the right instead of the left primary motor cortex. Fast spindles have shown hemispheric asymmetries, and motor-related spindle activity can be regionally specific, dominant over the hemisphere contralateral to the hand in use (Bódizs et al., 2017; Nishida & Walker, 2007). No motor tasks were performed before sleep, and participants were instructed to refrain from strenuous exercise before their visit, therefore spindle density may have been comparably lower in the stimulated right hemisphere, as it was contralateral to the non-dominant hand in our participants. Since the recording and stimulation configurations were mirrored across homologous centroparietal regions, we however expected the relevant relationship between spindle detection and stimulation to be preserved. (ii) Since we did not only apply spTMS to each condition but also ppTMS to quantify SICI, the conditioned ppTMS trials were pseudo-randomly interleaved with spTMS trials within each acquisition block. (iii) To save valuable sleep time given the need to collect sufficient trials across ten conditions, and as ITI distributions were not expected to differ across conditions owing to the pseudorandom counterbalancing within blocks, we did not include dummy pulses if no spindle was detected after a maximal inter trial interval of 7 s had passed. This could have prevented extreme ITI durations from influencing individual MEP amplitudes, although no bias towards any particular phase condition would be expected (Hassanzahraee et al., 2019). (iv) While our starting intensities for the spTMS trials were calculated as 120% and not 130% RMT, raw peak-to-peak spTMS MEP amplitudes across spindle-phase conditions were comparable (450-550 µV in our vs 400-500 µV in the previous study), ruling out differential floor effects. We had to increase stimulation intensities during sleep based on recalibrated RMTs in 12 of 20 participants, in whom MEP amplitudes especially of the ppTMS conditions were consistently < 50 µV. Intensity changes were accounted for in the subsequent analysis. (v) Instead of the arithmetic mean, we took the geometric mean of the MEP amplitude distribution of each condition, since amplitude distributions were skewed, as is typically the case with any measurements with a lower bound of zero. The geometric mean is equivalent to log-transforming MEPs, as is generally recommended, taking the arithmetic mean, and exponentiating, i.e. back-transforming the arithmetic mean to the original µV scale for easier interpretation than log-µV values. (vi) Lastly, our spindle-free baseline was implemented as being time-locked to just over 2 s post-spindle detection, which, depending on the definition may still lie within the spindle refractory period. However, this should not explain the difference in results since the previous study did not find a significant difference in CSE between the true spindle-free stimulation condition and the refractory period condition, which was implemented as 1.5 s after the spindle-centre (Hassan et al., 2025), and we replicated the general suppression of CSE during spindles relative to our spindle-free baseline.

### No modulation of SICI by sleep spindles

Contrary to our expectations, we did not find the spindle-related reduction in CSE to be mirrored by an increase in SICI. In light of the falling-phase driven pulsed suppression of excitability in the previous study (Hassan et al., 2025), and given the strong links between spindles and inhibition in animal work (Niethard et al., 2018; Peyrache & Seibt, 2020; Seibt et al., 2017), we had expected the reduction in excitability to be accompanied by a modulation in SICI. While we did not find a phase-specific modulation of excitability, the absence of an increase in SICI during spindles relative to spindle-free periods was unexpected.

These results may suggest that GABA-A receptor mediated intracortical inhibition may not the mechanism driving the reduction in CSE. Potentially other mechanisms like glutamatergic SICF or GABA-B receptor mediated inhibition might be involved. It is important to note that SICI was still consistently expressed during each spindle phase, and did not disappear like ICF has been found to do during REM sleep. To date, only on other study has applied real-time EEG-triggered ppTMS to quantify changes in SICI across oscillatory phase, namely of the motor cortical mu-alpha oscillation, which overlaps in frequency with spindles (Bergmann et al., 2019). They too found that a modulation by CSE, specifically a phase-dependent facilitation, was not accompanied by a modulation in SICI. They argued that in an environment of stable CSE, changes in SICI should be measured either following a change in the excitability of SICI-mediating inhibitory interneurons, for example PV+ cells, or following a change in the efficacy of their GABAergic transmission (Bergmann et al., 2019; Ilić et al., 2002). Yet in the context of a phasic modulation of excitability, a stable level of SICI, calculated as a relative measure of % change from TS MEPs, implies a covarying change in the absolute inhibition level. In our case, this implies that reductions in excitability would have been met with proportional attenuation of inhibition to maintain a stable SICI ratio across spindle phases. This would mean that the excitability of both pyramidal cells and the specific inhibitory interneurons contributing to the GABA-A receptor mediated SICI were proportionally modulated. A reduction in the net glutamatergic excitatory drive for example, could have led to the reduction in MEP amplitudes, with inhibitory interneurons receiving similarly lower levels of activation and consequently expressing proportionately less net inhibition, resulting in a maintenance of the SICI ratio.

Since a reduction in excitability was found at the corticospinal level, it could in theory be possible that any present inhibition is modulated at the spinal and not the cortical level. This is however not supported by the spindle literature. In fact, as detailed in the previous section, spindles have been closely associated with inhibition through their intricate coordination of cortical interneurons (SOM-In↓, PV-In↑), which promotes a local dendritic influx of Ca2+ that facilitates Ca2+-dependent plasticity processes including CaMKII activation (Fink & Meyer, 2002; Lüthi, 2014; Peyrache & Seibt, 2020).

A potential alternative explanation for our results is that floor effects precluded us from estimating the true levels of SICI. Since TS MEP amplitudes nonlinearly bias SICI estimates, the reduction in MEP amplitudes during all spindle-phases relative to spindle-free periods of N2/N3 sleep may have led to an underestimation of the relative strength of SICI (Cerins et al., 2022; Cirillo & Byblow, 2016; Higashihara et al., 2020). For this reason, threshold tracking approaches to SICI (T-SICI) have become more prevalent, in which the TMS intensity is continually adjusted over time to maintain a constant target MEP amplitude, and required intensity adjustments are recorded as the dependent variable. Unfortunately, this approach is not feasible during sleep, although an adjustment for excitability drifts across time would be highly desirable across different sleep stages, oscillations and phases. This is primarily because T-SICI requires significantly higher stimulation intensities than the standard approach, which would lead to more frequent TMS-induced arousals and exclusion of participants. Since it already proved difficult to recruit good supine sleepers who did not wake from stimulation during sleep, the likelihood of a participant being included in the study also depended on their RMT. At the end of each session of a pilot study, we increased spTMS intensities until they woke participants up, and found that most participants did not tolerate stimulation intensities during sleep higher than 80-85% MSO with our MagVenture MagPro X100 with MagOption stimulator.

### Preliminary evidence for a modulation of SICI in addition to excitability by SO

Our exploratory analyses of SO-presence on CSE and SICI during sleep spindles revealed that SOs not only increased excitability, but also significantly reduced SICI relative to isolated spindles.

The first finding is consistent with the previous finding of the SO up-state being more excitable than the down-state (Bergmann et al., 2012). Firstly, fast spindles, which were targeted in this study, tend to nest in the depolarising SO up-state, so this is the phase expected to have been predominantly targeted in our data (Mölle et al., 2002, 2011; Staresina et al., 2015). Secondly, SOs were detected post-hoc across multiple C4 area channels by their negative peaks within pre-TMS windows of 1.5s. SOs could not be detected using typical morphological, duration, fixed amplitude or zero-crossing criteria, as their EEG-trace was prematurely cut off at varying timepoints by the TMS-pulse. This means that negative peaks had to already have exceeded the detection threshold, reducing the likelihood that the TMS pulse coincided with the initial falling phase, but rather with the rising phase or up-state of detected SO. The limited detection window length ensured that the SO was still ongoing at the time of stimulation. Nevertheless, we cannot reliably verify the exact predominantly targeted phase of SOs at the timepoint of stimulation was the up-state, as we do not know the signal’s trajectory past the timepoint of stimulation. Subsequent EEG-data is not informative as it is contaminated by the high-amplitude TMS artifact and the TMS-evoked potential, or as is the case when stimulating during sleep, a TMS-induced SO (Massimini et al., 2007).

Since the previous study on excitability across SO-phases did not include a SO-free baseline condition, it remained unclear whether the modulations by phase also reflected symmetric net increases and decreases in excitability relative to SO-free N2/N3 sleep (Bergmann et al., 2012). We included two exploratory tests of the effect of SO-presence on excitability and SICI in the absence of spindles, i.e. including only spindle-free trials. Surprisingly they revealed the same pattern of SO-presence relatively increasing excitability whist reducing SICI, even though SOs may have been sampled at any phase angle in the absence of a spindle at stimulation. In relation to the previous investigation of SO-phase on excitability, these results suggest that at least the level of excitability during the SO up-state, if not the level of excitability across the entire SO, might be higher than that during SO-free N2/N3 sleep.

Together, these results offer new insights into the interplay between SO and spindles, which had previously only been investigated in isolation of one another, but need to be treated as preliminary since only ∼21% of all collected trials contained SOs leading to lower trial numbers in these analyses. One should also note that any SO-free trials were more likely to occur during N2 sleep whilst those with SOs present were more likely to have occurred during N3 sleep. Since the literature is mixed regarding which stage shows the lowest excitability (Bertini et al., 2004; Grosse et al., 2002; Salih et al., 2005), it remains unclear whether the biased occurrence of trials in N2 vs N3 in our SO analyses might be affected by baseline changes in excitability across sleep stage, or not. In contrast to our more ambivalent findings regarding SICI during spindles, the SO-related reduction in SICI cannot be attributed to the known positive relationship between TS MEP amplitude and SICI (Cerins et al., 2022; Cirillo & Byblow, 2016; Higashihara et al., 2020). With a relative increase in excitability during the SO up-state, this confound would predict increased instead of reduced SICI, ruling it out as alternative explanation. Nonetheless, the findings of a modulation of excitability and SICI by SO presence, or more likely specifically by the SO up-state, require replication via a dedicated real-time SO-phase triggered TMS study assessing both excitability and SICI across SO phases, ideally contrasting isolated SOs with SOs nesting spindles, and including a SO-free N2/N3 sleep baseline.

### Modulation of excitability and SICI across vigilance states

In line with previous studies, we found a strong suppression of CSE from pre-sleep wakefulness to both spindle-free N2/N3 sleep and to sleep spindles (averaged across phase). For all wake-sleep analyses, only blocks of data acquired at the same stimulation intensities as during preceding wakefulness were included to ensure a fair comparison. This reduced the sample sizes to 18 and 14 participants for spindle-average and spindle-free comparisons, respectively, as in 12 of 20 participants’ stimulation intensities had to be adjusted early in the night due to particularly strong suppression of MEP amplitudes below an average of 50 µV, leaving several participants with no or insufficient sleep data to be included in these analyses.

Previous findings regarding SICI had been less conclusive, with one study reporting a relative increase in SICI from wakefulness to N3/N4, but not to N2, and another study finding a significant increase from wakefulness to a condition of mixed N2/N3 sleep, in which N2 sleep had dominated (Avesani et al., 2008; Salih et al., 2005). In line with these studies, we found a non-significant trend (*p* = .052) towards an increase in SICI from preceding wakefulness to sleep spindles, and a marginally significant increase to spindle-free N2/N3 sleep. As the proportion of N2 sleep outweighed that of N3 sleep in our data, these borderline results point towards a likely discrepancy in the amount of inhibition present across N2 and N3 sleep. A comprehensive investigation of the changes in CSE, SICI and SICF across wakefulness, N1, N2, N3 and REM sleep with sufficient trial numbers in each condition, and ideally during periods of SO and spindle-free sleep, is still lacking. Based on our CSE and previous CSE and SICI results, SICI might be stronger during N3 than N2 sleep, whereas our SO results indicate the SICI to be reduced during SO, which are predominant during N3.

Any increases in SICI in the presence of smaller TS MEP amplitudes due to reduced CSE, as is the case during any wake-sleep comparisons, cannot be attributed to the confounding positive relation between MEP amplitudes and SICI (Higashihara et al., 2020). Instead, SICI is more likely underestimated because of floor effects. Therefore, our findings indicate that the strong suppression in CSE from wakefulness to N2/N3 sleep and sleep spindles is at least partially mediated by GABAergic intracortical inhibition mechanisms, and cannot be solely driven by reductions in glutamatergic signalling or mechanisms at the spinal level.

### Limitations

Several methodological considerations should be taken into account when interpreting the present findings. Real-time EEG-triggered TMS during sleep poses an inherent trade-off between maintaining stable and precisely targeted stimulation and preserving natural sleep. To ensure consistent coil positioning on target, participants slept in supine position with their head stabilised using a vacuum pillow, and the TMS coil secured by a mechanical arm. Although recordings were conducted in a dedicated sleep cabin using a conventional bed to maximise comfort, this constrained sleeping position limited our recruitment to individuals able to sleep reliably under these conditions. It also limited recording duration, requiring data to be pooled across nights in 14 of 20 participants. The use of multiple shorter nocturnal recordings represented a pragmatic compromise that prioritised stimulation accuracy and sleep quality. Future studies may benefit from robot-navigated coil positioning, which could compensate for head movements while permitting less restricted and non-supine sleeping positions, and in effect longer sleep recordings.

A second consideration concerns the definition of the spindle-free baseline N2/N3 sleep condition. Stimulation was delivered 2 s after spindle detection to target periods with a low spindle probability, while maintaining a comparable N2/N3 sleep state. This provided a practical control during comparably deep sleep, but did not constitute a fully independent spindle-free condition, and may have overlapped with the spindle refractory period. Trials containing newly emerging spindle activity were removed post-hoc, slightly reducing trial numbers in this condition. Although previous work had found no significant difference in CSE between spindle-refractory and spindle-free periods, the absence of an unambiguous spindle-free condition in our study, which may have overlapped with the refractory period, highlights the importance of including a spindle-independent spindle-free condition in future work (Hassan et al., 2025). Similarly, SOs were identified post-hoc rather than incorporated within the real-time detection algorithm. Approximately 21% of spindle trials coincided with an SO, and were therefore excluded from the primary analyses of isolated spindles. While pooling across spindle phases provided sufficient trials for a secondary comparison of SO-nested and isolated spindles, the study was not specifically designed to investigate interactions between SO and spindle phase. In future, using real-time algorithms that jointly detect SOs and spindles would enable these events to be experimentally dissociated, and their relationship to be systematically explored.

A further limitation concerns the choice and adjustment of TMS stimulation intensities. Stimulation intensities were initially intended to be individualised based on pre-sleep input– output curves. Yet, obtaining reliable curves proved impractical immediately before nocturnal naps, because the extended calibration procedure interfered with participants’ ability to maintain stable wakefulness and produced variable estimates. Therefore, RMT-based intensities were considered to be more time-efficient and reliable, especially as they had yielded comparable stimulation intensities in participants with usable input–output curves during piloting. Furthermore, the marked reduction in MEP amplitudes from wakefulness to sleep and during sleep spindles presents a particular challenge for measuring SICI, as SICI depends on the amplitude of the unconditioned test response (Cirillo & Byblow, 2016). Stimulation intensity therefore required adjustment during sleep in 12 of 20 participants, to minimise floor effects and maintain measurable MEPs, especially in the ppTMS conditions. Whilst necessary, this approach does not eliminate potential bias arising from differences in TS MEP amplitude between conditions, and may have reduced the sensitivity to modulations of SICI. Calibrating test stimulation to 1mV and not 120% RMT before sleep, or using threshold-tracking approaches (T-SICI), may provide more comparable TS MEP amplitudes across brain-states. However, such approaches require higher stimulation intensities, which increase the strength of auditory and somatosensory TMS co-stimulation, and consequently the likelihood of sleep disruption and drop-outs. In the present study, stimulation parameters therefore represented a deliberate compromise between optimising the physiological TMS measure and maintaining stable sleep.

Finally, SICI captures only one mechanism within specific intracortical circuitry which could contribute to the spindle-related modulation of CSE. The absence of spindle-related changes in SICI therefore does not exclude contributions from other inhibitory or facilitatory mechanisms. Complementary ppTMS protols like SICF could provide a more comprehensive characterisation of spindle-related cortical network dynamics. Although SICF was considered during piloting, its inclusion would have increased the number of conditions from 10 to 15. We prioritised reliable trial numbers across fewer conditions. Future studies combining improved real-time detection with longer and less constrained recordings using a collaborative robot, could characterise the modulation of several other local network properties across spindle phase.

Overall, our methodological choices reflected the competing requirements of precise spindle phase-targeted TMS, reliable physiological EEG, PSG and EMG measurements, and preservation of high-quality sleep. While they constrain the interpretation of some findings, the present approach enabled repeated measurement of CSE and SICI at precisely targeted spindle phases during human NREM sleep. Further methodological advances in real-time event detection and the implementation of automated coil positioning should allow future studies to build on our approach while reducing constraints.

### Outlook

Our findings provide several ideas for follow-up studies. During wakefulness, the mu-alpha rhythm has been well-characterised in terms of its oscillatory profile of CSE and inhibition (Bergmann et al., 2019; Schaworonkow et al., 2019; Stefanou et al., 2018; Wischnewski et al., 2022; Zrenner et al., 2018, 2023), and has more recently been shown to demonstrate phase-dependent plasticity. Repeatedly applying high-frequency rTMS bursts only induced long-term potentiation LTP-like increases in CSE when applied to the most excitable mu-phase, the trough, but not when targeting the least excitable mu-peak, whereas random-phase targeting resulted the induction of lasting long-term depression (LTP)-like decreases in excitability (Baur et al., 2020, 2022; Zrenner et al., 2018). Given the hierarchical nesting of SOs, spindles and ripples coordinating the induction of plasticity underling memory consolidation during sleep, a similar modulation of the effectivity and or directionality of long-term plasticity may be found for SOs but more specifically, spindles. Whilst we did not find a significant modulation of CSE by any one spindle phase, the pattern of CSE across phases was generally in line with previous results as well as theoretical expectation based on spindle-ripple nesting, with the spindle falling flank and phase showing the lowest and the peak the highest levels of excitability (Hassan et al., 2025; Staresina et al., 2015). This suggests that these phases might reflect the optimal timepoint most receptive to the induction of plasticity. Future real-time EEG-triggered TMS studies investigating spindle-phase dependent plasticity by repetitively delivering rTMS bursts to the same spindle phase could consequently compare the effectivity of plasticity induction during the spindle falling phase or trough, to the spindle peak. Since spindles have been associated with both LTP- and LTD-like plasticity (Bergmann et al., 2008; Fernandez & Lüthi, 2020; Rosanova & Ulrich, 2005), LTD-plasticity inducing rTMS protocols may be more effective when time-locked to the spindle falling phase or trough, while LTP-like plasticity may be facilitated by the spindle peak.

Next, a comprehensive characterisation of CSE, SICI and SICF across vigilance and all AASM-defined sleep stages (Iber et al., 2007) would set the ground in terms of quantifying baseline drifts in cortical and corticospinal activity in the human cortex over longer periods of time. With sufficient trial numbers in each condition, and stimulation only occurring during SO- and spindle-free sleep, such a study could help clarify existing ambiguities for example regarding changes with deepening NREM sleep.

Furthermore, our findings point towards a potential modulation not only of excitability but also of SICI across SO phases, which could be probed using real-time SO-phase-triggered TMS of at least four oscillatory phases for a more fine-grained resolution, and should include a SO-free baseline of N2/N3 sleep (Bergmann et al., 2019). Given their hierarchical coordination with sleep spindles, it is important to better understand this large-scale modulation which sets the scene for spindles and ripples. Ideally, such an investigation would differentiate between isolated SOs and SO-spindle complexes.

## Conclusion

This study provides the first characterisation of local GABA-A receptor-mediated SICI across sleep spindle presence and phase in the human cortex. We broadly replicated the previously reported suppression of corticospinal excitability during NREM sleep, and in particular during spindles, with the numerically lowest excitability observed during the falling flank and trough, although no significant phase-specific modulation was observed. In contrast, we found no evidence that SICI was modulated by either spindle presence or phase. SICI, generally increased during NREM sleep, was however transiently reduced during the occurrence of SOs, both with and without concurrent spindles. This finding suggests that intracortical disinhibition may contribute to the established increase in excitability during the SO up-state relative to the down-state, contributing to its role in memory consolidation and synaptic rescaling. Together, our findings indicate distinct contributions of spindles and SOs to cortical network dynamics and provide new insight into the transient modulation of excitability and inhibition during human NREM sleep.

## Acknowledgements

AI (GPT-5.6 Sol and Consensus) was used to help refine the writing style and reduce word count across several sections of this paper, after which these sections were again reviewed and edited by the authors, who take full responsibility for the content.

